# Nitrogen niche partitioning between tropical legumes and grasses conditionally weakens under elevated CO_2_

**DOI:** 10.1101/2023.01.16.524162

**Authors:** Amber C. Churchill, Haiyang Zhang, Gil Won Kim, Karen L. M. Catunda, Ian C. Anderson, Forest Isbell, Ben Moore, Elise Pendall, Jonathan M. Plett, Jeff R. Powell, Sally A. Power

## Abstract

Plant community biodiversity can be maintained, at least partially, by shifts in species interactions between facilitation and competition for resources as environmental conditions change. These interactions also drive ecosystem functioning, including productivity, and can promote over-yielding-an ecosystem service prioritized in agro-ecosystems, such as pastures, that occurs when multiple species together are more productive than the component species alone. Importantly, species interactions that can result in over-yielding may shift in response to rising CO_2_ concentrations and changes in resource availability, and the consequences these shifts have on production is uncertain especially in the context of tropical mixed-species grasslands.
We examined the relative performance of two species pairs of tropical pasture grasses and legumes growing in monoculture and mixtures in a glasshouse experiment manipulating CO_2_. We investigated how over-yielding can arise from nitrogen (N) niche partitioning and biotic facilitation using stable isotopes to differentiate soil N from biological N fixation (BNF) within N acquisition into aboveground biomass for these two-species mixtures.
We found that N niche partitioning in species-level use of soil N vs. BNF drove species interactions in mixtures. Importantly partitioning and overyielding were generally reduced under elevated CO_2_. However, this finding was mixture-dependent based on biomass of dominant species in mixtures and the strength of selection effects for the dominant species.
This study demonstrates that rising atmospheric CO_2_ may alter niche partitioning between co-occurring species, with negative implications for the over-yielding benefits predicted for legume-grass mixtures in working landscapes with tropical species. Furthermore, these changes in inter-species interactions may have consequences for grassland composition that are not yet considered in larger-scale projections for impacts of climate change and species distributions.

**Graphical abstract:** (Image by H. Zhang): Among our tropical pasture species we found that grasses (dotted lines) grown in monoculture rely fully on soil nitrogen (N), while legumes (solid lines) grown in monoculture relied approximately equally on soil N and biological nitrogen fixation (BNF) to meet N requirements. When grown with tropical grasses, however, legumes shifted to rely more strongly on BNF, indicative of niche partitioning and decreased competition for soil nutrients with grasses. This separation of niche space was weakened under elevated CO_2_ conditions, ultimately reducing legume production.

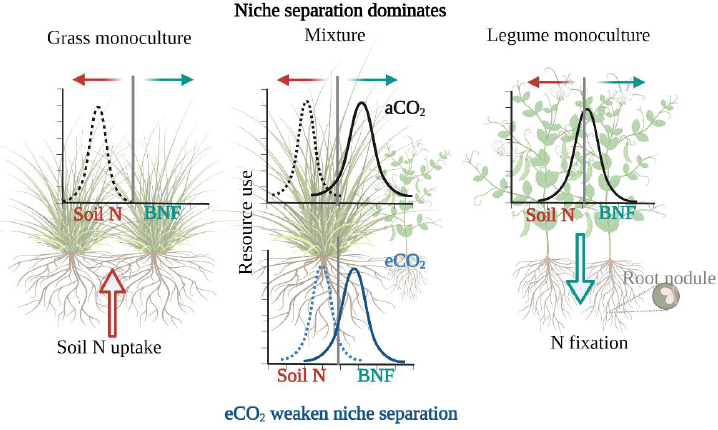

## Introduction

Interactions between plant species within communities and ecosystems vary over time and as environmental conditions change, thereby driving shifts between competition and facilitation for resources (Adler et al., 2012; Arfin Khan et al., 2014). Niche partitioning is a mechanism allowing the co-existence of numerous species that may overlap in fundamental niche space but differ in realized niche space to reduce impacts from competition (Jumpponen et al., 2002; McKane et al., 2002; Zuppinger-Dingley et al., 2014). Additionally, changes in niche space associated with interspecific interactions are a common feature of net biodiversity effects whereby interactions among community members result in greater ecosystem functioning, relative to the sum of individual contributions (Barry et al., 2019; Loreau et al., 2008; Loreau & Hector, 2001); when applied to productivity this concept is often referred to as over-yielding (Vandermeer, 1981). Quantifying mechanisms underpinning interspecific interactions, such as niche partitioning, is fundamental to understanding relationships between biodiversity and ecosystem function.

Nitrogen (N) is a primary often-limiting nutrient that plays an important role in structuring community interactions and ecosystem function alongside phosphorus (P) and other mineral nutrients (Fay et al., 2015; Vitousek & Howarth, 1991). Plants that host N-fixing bacteria (i.e. N-fixers), such as legumes, are often considered to have an advantage under low soil N conditions as the mutualism can provide an alternative route for accessing N. This biological N fixation (BNF), therefore, can promote biodiversity and ecosystem function, via N niche partitioning allowing different community members to rely on alternative sources of N or facilitation where N resources from BNF are shared among community members (Barry et al., 2019; Elias & Agrawal, 2021; Lee et al., 2003; Mulder et al., 2002; Perring et al., 2010). These processes contribute to the Net Biodiversity Effect through complementarity effects, where more diverse communities contribute to ecosystem functions greater than the weighted average of constituent monoculture species, and selection effects where species with greater than the average among-species-yields dominate in mixture.

In communities containing legumes, BNF is often assumed to result in N facilitation and an increase in the overall pool of ecosystem N available for neighbouring plants (Lee et al., 2003) that promote and subsequently increase production among component species (Barry et al., 2019; Fornara & Tilman, 2008; von Felton et al., 2009). There are two main pathways enabling this outcome: 1) direct acquisition of BNF resources from legumes by non-legumes (facilitation; i.e. grasses) and 2) via niche partitioning where grasses compete less with legumes for shared soil nitrogen, given that legumes have access to a non-soil pool of N (Barry *et al*., 2019). There are a number of biotic feedback-based mechanisms enabling the sharing of BNF derived N to non-legume community members, including decomposition of legume leaf litter (Makkonen et al., 2012; Wei et al., 2019), or roots (Fustec et al., 2009; Mclaren & Turkington, 2010) that lead to higher soil N. Additionally, some literature has highlighted the potential importance of root exudation of high N-containing compounds that impact N (or carbon: C) availability in the rhizosphere (Henneron et al., 2020; Paynel et al., 2001), or transfer through the activity of soil fauna (Gilarte et al., 2020), thereby promoting increased N availability for surrounding community members. Alternatively, niche partitioning can support greater overall ecosystem N availability as legumes rely on BNF for N resources, leaving the soil N pool available to neighbouring community members (Ashworth et al., 2018). These pathways offer numerous strategies that ultimately promote increased community productivity in the presence of legumes representing a potential for over-yielding.

Atmospheric CO_2_ concentrations have increased by 50% since the industrial revolution (IPCC, 2022). There are many ecosystem consequences associated with this change, including an increase in plant productivity (de Graaff et al., 2006; Gill et al., 2002), altered plant community composition and diversity (Reich, 2009; Zelikova et al., 2014), and modified plant interactions within a community that impact ecosystem scale processes (Blumenthal et al., 2013; Gonçalves de Oliveira et al., 2021; Maschler et al., 2022). However, some of these patterns may only occur when water is limiting, nutrients are abundant (Reich et al., 2014; Rogers et al., 2009; Soussana & Lüscher, 2007), or when plant diversity is high (Reich et al., 2001; Shaw et al., 2002). Under well-watered conditions, eCO_2_ generally benefits the C_3_ photosynthetic pathway over the C_4_ pathway (used by warm season or tropical grasses) due to differences in carbon capture (Ainsworth & Long, 2005; Reich *et al*., 2014, but see Reich *et al*., 2018). Furthermore, eCO_2_ is reported to increase belowground biomass, especially for fine roots (Nie et al., 2013), which may increase access to soil nutrients (de Graaff et al., 2006; Dieleman et al., 2012; Reich et al., 2001).

Many studies have found that eCO_2_ fertilization effects are larger for N fixers than grasses due to greater C resource availability to support the energetically demanding process of N-fixation (Liang et al., 2016). However, N-fixer responses in natural ecosystems have been varied, as potential benefits from increased photosynthesis can become constrained by other limiting nutrients such as phosphorus (Coventry et al., 1985; Hungate et al., 2004; Rogers et al., 2009; Soussana & Lüscher, 2007), or reduced due to temperature or drought conditions (Rogers et al., 2009). Even so, most studies find decreased foliar N concentrations for non-N-fixers under eCO_2_ with subsequent increases in C:N ratios in plant tissues (Ainsworth & Long, 2005; Augustine et al., 2018; Gojon et al., 2022; Rogers et al., 2009). However, within mixed grass-legume grassland communities, leaf N and C:N ratios are less likely to be impacted by eCO_2_ for the legume component of the sward (Rogers et al., 2009; Winkler & Herbst, 2004). These findings suggest that grassland responses to eCO_2_ may depend on the presence of legumes (Jones & Donnelly, 2004). Importantly, the perceived ‘benefit’ of eCO_2_ for tropical and subtropical ecosystems is less well documented, as most studies have focused on temperate species, with C_3_ photosynthetic pathways (Naudts et al., 2013). The potential impacts of eCO_2_ on nutrient dynamics in C_4_ grasses, including those in mixed C_3_-C_4_ systems is of particular interest, due to the increasing abundance of warm season grasses in many areas of the world (McKeon et al., 2009). Given the diverse predicted outcomes for ecosystems as eCO_2_ concentrations increase (Cannell & Thornley, 1998; de Graaff et al., 2006; Dijkstra et al., 2010), there is a need for improved understanding of the impacts of rising CO_2_ in managed and natural grasslands. These include ongoing shifts in grassland composition, driven by species differences in response to climate change (McKeon et al., 2009; Munson & Long, 2017; Sherry et al., 2007; C. Wang et al., 2013; Xie et al., 2022; Zelikova et al., 2014). Increased understanding of the processes that underlie plant-plant interactions between tropical grasses and legumes will therefore increase the accuracy of predictions for how grassland species composition and productivity will be impacted as atmospheric CO_2_ concentrations continue to rise and human-induced shifts in ecosystem biodiversity are realized.

To address this area of research, we examined potential mechanisms by which eCO_2_ might benefit warm season (C_4_) grasses grown in mixed swards with tropical legumes. Specifically, we evaluated the potential for increased production and nutrient availability within two-species swards relative to individual species, driven by biologically fixed N (from rhizobia) supported by legumes and accessed by grasses. We hypothesized that 1) grasses grown with legumes benefit in terms of both nutrition and productivity, relative to those growing in monoculture, due to access to BNF from legume rhizobia; 2) tropical legumes grown under eCO_2_ will increase production as a consequence of increased carbon-use-efficiency; 3) increased C availability under eCO_2_ will increase legume N fixation and both legume shoot N and soil N content; and 4) increased available N due to BNF will promote overyielding and a net biodiversity effect in two-species mixtures driven by a combination of complementarity and selection effects that are enhanced by eCO_2_.

## Materials and Methods

### Experimental design

This experiment was conducted in a glasshouse at Western Sydney University’s Hawkesbury Campus, Richmond, New South Wales (33°61’S,150°74’E), Australia. We used six separate glasshouse chambers, with three randomly assigned to ambient conditions (410 - 430 ppm; aCO_2_) and three an elevated CO_2_ regime (630 - 650 ppm; eCO_2_). The ambient treatment matches contemporary atmospheric concentration, while the elevated regime was consistent with the upper-end of predicted end-of-century increase in global atmospheric [CO_2_] (Szulejko et al., 2017). We maintained a consistent temperature regime across all chambers at a daily average of 27°C (**Fig S1A**). This included a temperature ramp to mimic diurnal patterns (00-06 hours: 22°C, 06-09 hrs: 27°C, 09-18 hrs: 32°C, 18-21 hrs: 27°C, 21-00 hrs: 22°C) for spring to summer temperatures for areas currently cultivating tropical pasture in northern NSW and southern Queensland (BOM, 2020). Additionally, all chambers were controlled at 50-60% relative humidity (**Fig S1B**) with natural lighting between 11 hours (August) and 14 hours (December) per day.

The soil used in this experiment was collected adjacent to the Pastures and Climate Extremes field experimental facility (Churchill et al., 2022). Soil was sieved to 5 mm before being air dried and mixed with quartz sand (soil:sand 7:3 v/v). Each pot (3.7L, 150 mm diameter, 240 mm height) was filled with 3.9 kg of the soil-sand mixture before planting (Zhang et al., 2021). The final soil mixture was ∼ 84% sand, with an average pH of 5.6, plant available N at 55 mg kg^−1^, plant available (bray) P at 39 mg kg^−1^ and soil organic carbon at 1.1%. All pots were maintained under well-watered conditions using an automated irrigation system such that soil water capacity was brought to 100% every two days (Zhang et al., 2021). In order to promote legume nodulation, no nutrients were added.

Two tropical grasses (C_4_) and two tropical legumes (C_3_) were selected, based on current widespread use or consideration for future use by key pasture industries within Australia (**Table 1**). All species are commonly grown in multi-species improved pastures in southern Queensland and northern New South Wales, Australia, as well as among other sub-tropical grasslands world-wide (Thomas & Sumberg, 1995; Bell *et al*., 2012; Miegoue *et al*., 2016; Melesse *et al*., 2017; DPI, 2018). Specific species-pairs were the legume *Macroptilium bracteatum* (Nees. & Marti.) growing with the grass *Chloris gayana* (Kunth), and the legume *Desmodium intortum* (Mill) Urb with grass *Panicum maximum* var. *trichoglume* (Robyns).

**Table 1.** Tropical species and associated mixtures.

| Scientific Name* | Common Name | Variety | Growth form | Paired species |
| --- | --- | --- | --- | --- |
| *Chloris gayana* (Kunth, 1830) | Rhodes grass | Katambora | Grass | Burgundy |
| *Desmodium intortum* (Mill. Urb.) | Greenleaf Desmodium | Greenleaf | Legume | Panic |
| *Macroptilium bracteatum* (Nees & Mart.) | Burgundy Bean | Presto | Legume | Rhodes |
| *Panicum maximum* var. *trichoglume* (Robyns) | Panic grass | Megamax 059 | Grass | Desmodium |

*\*Species are referenced by genera in text*

Specific species pairing within pots was randomly assigned at project initiation, as all species are typically grown in mixtures. We included non-factorial pairings between grasses and legumes to maximize species included in the study, while balancing time and space constraints in the greenhouse. All seeds were supplied by Heritage Seeds and contained AgriCote advanced seed coating technology; legume seed came pre-inoculated with appropriate rhizobia (Barenbrug Co., Heritage Seeds, New South Wales, Australia). Seeds were planted directly into seedling trays (∼5 seeds per cell) and thinned to a single individual over time. After three weeks, two individuals for the monoculture or one individual from each species for the mixture were then transplanted into the pots. The experimental design therefore included the two CO_2_ treatments (aCO_2_ and eCO_2_), six types of plant combinations in pots (grass_1_, legume_1_, mixed_1_, grass_2_, legume_2_, mixed_2_), and twelve replicates per plant combination with four of these in each of three chamber replicates per CO_2_ treatment). This included a total of 288 pots distributed among six climatically isolated greenhouse chambers.

### Aboveground sampling

After transplanting the seedlings, all plants were grown for 11 weeks before a final harvest, just as individuals were reaching the flowering phenophase. All aboveground standing plant material (shoot; leaves and stems/tillers together) was cut at soil level and for mixed species pots, separated by species, then weighed. To accommodate differing methodologies for nutritional analysis of harvested biomass (Catunda et al., 2022), the fresh aboveground biomass was frozen at -18^°^C. Within one month all samples were then microwaved at medium power for 90 seconds to halt enzymic degradation of plant tissue compounds (Landhäusser et al., 2018), and oven dried at 70^°^C for 48 hours then weighed. All dried plant material was homogenized, ground to pass through a 2-mm screen in a laboratory cyclone mill (Foss Cyclotec Mill, Denmark) then ball-milled to a fine powder (Retsch® MM400; Hann, Germany) for nitrogen (N) and isotopic (i.e. δ^15^N) measurements, which were determined using an elemental analyzer interfaced to a continuous flow isotope ratio mass spectrometer at the UC Davis Stable Isotope Facility (Davis, CA, USA).

### Belowground biomass

After aboveground material was removed, belowground biomass was determined. All pot soils were carefully sieved using 2 mm sieves to retain root fragments and roots were oven dried to constant mass. We assessed legume nodule number and weight for a subset of fresh root material in each legume- or mixed species pot by randomly subsetting roots and nodules from different locations within the root system (∼25g fresh root or ∼ 80% of total root mass). From this subset, we separated the nodules from the roots and weighed the fresh biomass for each component separately. Fresh nodules were incubated with acetylene for 1 hour and ethylene evolution tested as a proxy to estimate the average instantaneous rate of nitrogenase activity of the bacteria within the nodules using acetylene reduction assays (ARA; Plett *et al*., 2016). After ARA measurement, root nodules were oven dried and weighed as above. Final estimates of biological nitrogen fixation (BNF) activity were calculated as the rate of ethylene production per gram of incubated nodule dry biomass.

### Soil nutrients

Nutrient availability was assessed using ion exchange resins (IONAC NM-60 H^+^/OH^-^ Form; J.T. Baker) for nitrate (NO_3_^-^), ammonia (NH_4_^+^) and phosphate (PO_4_^-3^). We enclosed 5 g of mixed bed resin in a 3 cm x 5 cm nylon bag (42 μm mesh) and placed one bag in each pot in the upper 10 cm of soil. Resin bags were collected during the biomass harvest, briefly rinsed with deionized water to remove attached soil and root particles before immediate storage at - 70 °C prior to extraction with 0.5 M HCl solution through a Whatman No. 1 paper (Carrillo et al., 2012; Dijkstra et al., 2010). Extractable NO_3_^-^, NH_4_^+^ and PO_4_^-3^ concentrations in the resin bag were determined on a SEAL AQ2 Discrete Analyser (SEAL Analytical Inc., USA).

### Quantifying contributions of biological nitrogen fixation

The contribution of BNF to plant total N was calculated as described in Chalk *et al*. (2016) and Bell *et al*. (2017), where lower δ ^15^N is indicative of a higher N-fixation amount. The minimum observed values for δ^15^N for both legume species in this experiment (*B*; *Macroptilium*: -2.7 ‰ under eCO_2_ in a mixed pot; *Desmodium*: -2.0 ‰ under aCO_2_ in a mixed pot) were used to represent when the maximum possible amount of N is derived from BNF in our experimental conditions. Note that while this could overestimate the absolute estimate of BNF, the general patterns among different treatments should remain unaffected (Chalk et al., 2016). Mean δ^15^N values for companion grass species in monoculture at each treatment level of CO_2_ were used as baselines for soil derived N (*Ref*; *Chloris*: 3.9 ‰, 3.7‰; *Panicum*: 4.9 ‰, 4.4 ‰; aCO_2_, eCO_2_ respectively), and we then calculated the relative contribution of BNF to plant aboveground N following Equation 1:

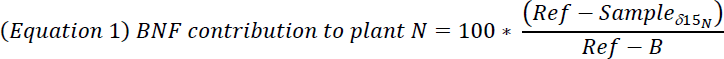

Calculations of the percentage of shoot N derived from BNF were performed for each species within a pot, based on species specific shoot δ^15^N. The contribution of soil N to shoot N was then calculated as the difference between 100% and BNF values for each species within a pot. Species level shoot N content derived from either BNF or soil was then calculated by multiplying total shoot N content by the relative percentage for both sources.

### Calculations and statistical analyses

Effects of CO_2_ treatment on aboveground plant responses measured on pot level totals (aboveground shoot biomass, total shoot N content, soil derived shoot N content, belowground biomass and resin extractable soil nutrients) were analysed using linear mixed effects models. For these analyses, ‘Species Pair’, ‘CO_2_’ and ‘Pot Type’ treatments were included as fixed effects with all two-way and three-way interactions. For analysing shoot % N we conducted a linear mixed effect model with ‘Species Pair’, ‘CO2’ and ‘Plant Type’. ‘Chamber’ was also included as a random effect in all models to account for non-independence among replicates within each chamber due to potential among-chamber environmental variation. Treatments of CO_2_ included two levels (aCO_2_ and eCO_2_), ‘Species Pair’ included ‘*Macroptilium* & *Chloris*’ and ‘*Desmodium* & *Panicum*’, the ‘Pot Type’ category included ‘Grass’, ‘Legume’, and ‘Mixed’ pots, and the ‘Plant Type’ category included ‘Grass’, ‘Legume’, grass grown with legume: ‘L-Grass’, and legume grown with grass: ‘G-Legume’. Nodule data were also analysed using linear mixed effects models, such that ‘CO_2_’, ‘Species Pair’, and ‘Pot Type’ were included as interactive fixed effects and ‘Chamber’ as a random effect. For these analyses ‘Pot Type’ only included ‘Legume’ and ‘Mixed’ pots.

Community over-yielding was determined as the increased productivity of a mixed pasture over the discrete contributions of the constituent species grown in monoculture (Isbell et al., 2018). We calculated over-yielding for each species pair based on mixture total shoot biomass (and total N content in aboveground plant material) and the associated shoot biomass of the constituent monoculture grass and legume species standardized by the number of individuals per pot in each chamber, following Equation 2:

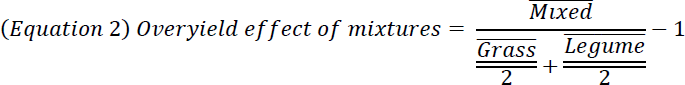

Effects of eCO_2_ on over-yielding were examined by calculating over-yield estimates based on average monoculture aboveground biomass by species and average mixed pots for the component species pairs in each chamber (n = 6; 3 for each CO_2_ level). To examine differences in over-yield effect between CO_2_ treatments and species pair swards, we then ran a linear mixed effects model with over-yield effect as the response, ‘CO_2_’ level and ‘Species Pair’ as the fixed effects and ‘Chamber’ as a random effect. Statistical significance of the effect size for over-yielding was determined by calculating mean treatment values and 95% confidence intervals (CI) across all replicates (n = 12 for each plant type and CO_2_ level) and over-yielding was deemed significant if CIs did not overlap with 0. For these results, CIs that did not overlap with 0 indicate either a gain in shoot biomass in mixed pots relative to monoculture pots (positive values) or a loss in shoot biomass in mixed pots relative to monoculture pots (negative values).

The net biodiversity effect, and its constituents-the complementarity effect and selection effect-were calculated following Loreau & Hector (2001) based on averages for monoculture and mixed pots of each species pair in each chamber and analysed statistically as described for the over-yield metric above. Net biodiversity effect was therefore defined as the difference between the observed yield of a mixture and the expected yield in the absence of complementarity or selection (Loreau & Hector, 2001). NBE is considered the net balance of complementarity effects and selection effects. Complementarity effects are positive when mixture yields are higher than expected based on the weighted average of constituent monoculture species, while selection effects are positive when species with greater than the average among-species-yields dominate in mixture.

All analyses were conducted in R version 4.1.1 (R Core Team, 2020) using the package lme4 (Bates et al., 2015) with statistical contrasts based on Kenward-Roger degrees of freedom calculated using the ‘Anova’ function from the ‘car’ package (Fox & Weisberg, 2019). Pairwise comparisons to determine treatment differences in plant responses for all models were conducted using the R package emmeans (Length, 2020) and using the Tukey method for *P*-value adjustment. All data used in these analyses are publicly available on Dryad (Churchill et al., 2024).

## Results

### Shoot biomass and N content

Shoot biomass differed strongly between our species pairs, with greater productivity in pots with the *Macroptilium* & *Chloris* monocultures and mixture, compared with *Desmodium* & *Panicum* (**Fig 1**; **Table 2**). Elevated CO_2_ (eCO_2_) had limited effects on shoot biomass, with increased productivity only observed for *Macroptilium* monoculture pots (**Fig 1a**). There were no significant productivity differences among pot types (grass, legume, mixed) for the *Macroptilium* & *Chloris* pairing, while *Desmodium* in monoculture produced less shoot biomass than *Panicum* or the mixed pot type (**Fig 1b**). Despite little change in pot-level biomass among monocultures and mixed pots, the contribution of grasses and legumes to mixed pot total biomass was heavily weighted toward grass production in both species-pairs, with *Chloris* contributing 65% and 61% of total pot biomass under aCO_2_ and eCO_2_ and *Panicum* 90% and 95%, respectively. Relative to growth of individual plants in monoculture pots, this change represented a 38%/52% gain in biomass under aCO_2_ and eCO_2_ by *Chloris* and 20%/28% reduction in growth by *Macroptilium* (**Fig 1a**). This trend was stronger for the *Desmodium* & *Panicum* species pair with 96%/87% gain for *Panicum* and 65%/86% loss for *Desmodium* between monoculture and mixed pots (**Fig 1b**).

**Figure 1.**
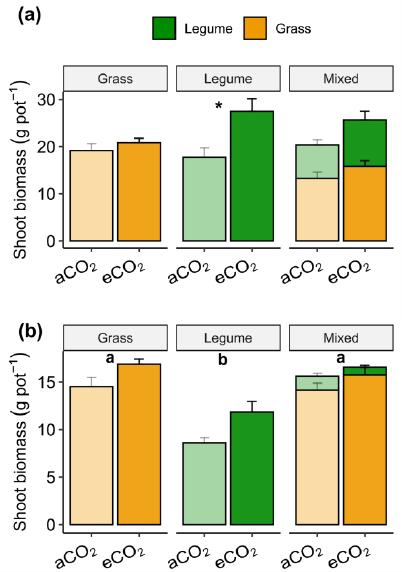
Shoot biomass of grass and legume grown as monoculture or mixtures for two species pairs under ambient (aCO_2_) and elevated CO_2_ (eCO_2_) comprising (a) *Macroptilium* & *Chloris* and (b) *Desmodium* & *Panicum*. Bars indicate mean values, plus standard error. Significant differences (p < 0.05) between CO_2_ treatments are indicated by ‘*’, and significant differences among pot types (grass, legume, mixed) are indicated by differing letter designations.

**Figure 2.**
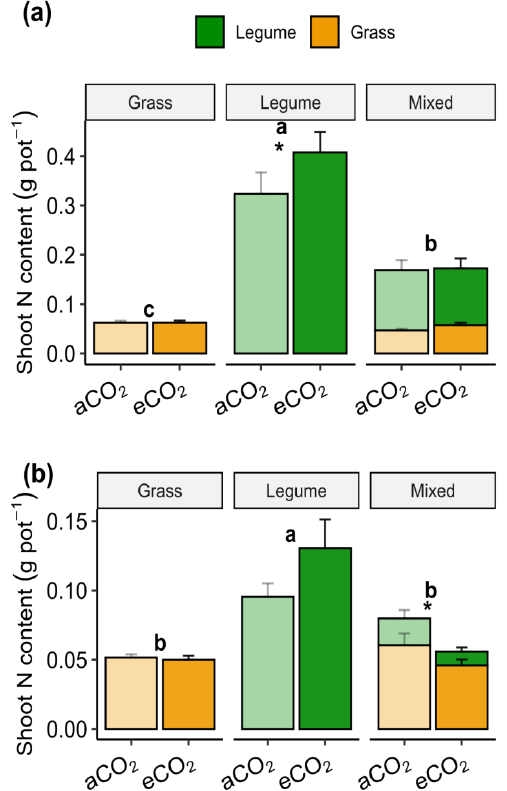
Shoot nitrogen (N) content (%N multiplied by shoot biomass) partitioned between grasses and legumes grown as monocultures or mixtures under ambient (aCO_2_) and elevated CO_2_ (eCO_2_), for (a) *Macroptilium* & *Chloris* and (b) *Desmodium* & *Panicum*. Bars indicate mean values, plus standard error. Notations follow Fig. 1.

**Table 2.** Effects of CO_2_ treatment (CO_2_; ambient, elevated), species pair (SP; *Macroptilium*/*Chloris* and *Desmodium*/*Panicum*) and pot type (PT; grass, legume, mixed-species), for pot level total shoot biomass, total shoot N, total shoot N derived from soil (shoot N-soil).

| Response | Factors | *F* Statistic* | *P* Value | *r*<sup>2</sup>-M | *r*<sup>2</sup>-C |
| --- | --- | --- | --- | --- | --- |
| Shoot biomass<sup>^</sup> | CO<sub>2</sub> | 5.23<sub>1,4</sub> | 0.08 | 0.40 | 0.45 |
|  | Species pair | 49.9<sub>1,143</sub> | <0.01 |  |  |
|  | Plant type | 12.1<sub>2,143</sub> | <0.01 |  |  |
|  | CO<sub>2</sub> x SP | 1.9<sub>1,143</sub> | 0.17 |  |  |
|  | CO<sub>2</sub> x PT | 2.5<sub>2,143</sub> | 0.08 |  |  |
|  | SP x PT | 5.6<sub>2,143</sub> | <0.01 |  |  |
|  | CO<sub>2</sub> x SP x PT | 1.0<sub>2,143</sub> | 0.38 |  |  |
| Shoot N content<sup>^</sup> | CO<sub>2</sub> | 0.3<sub>1,4</sub> | 0.60 | 0.65 | 0.66 |
|  | Species pair | 102.7<sub>1,143</sub> | <0.01 |  |  |
|  | Plant type | 79.4<sub>2,143</sub> | <0.01 |  |  |
|  | CO<sub>2</sub> x SP | 1.9<sub>1,143</sub> | 0.18 |  |  |
|  | CO<sub>2</sub> x PT | 3.7<sub>2,143</sub> | 0.03 |  |  |
|  | SP x PT | 14.7<sub>2,143</sub> | <0.01 |  |  |
|  | CO<sub>2</sub> x SP x PT | 0.4<sub>2,143</sub> | 0.64 |  |  |
| Shoot N-soil | CO<sub>2</sub> | 0.1<sub>1,4</sub> | 0.79 | 0.14 | 0.15 |
|  | Species pair | 17.8<sub>1,143</sub> | <0.01 |  |  |
|  | Plant type | 2.7<sub>2,143</sub> | 0.07 |  |  |
|  | CO<sub>2</sub> x SP | 0.4<sub>1,143</sub> | 0.54 |  |  |
|  | CO<sub>2</sub> x PT | 0.1<sub>1,143</sub> | 0.92 |  |  |
|  | SP x PT | 0.04<sub>2,143</sub> | 0.95 |  |  |
|  | CO<sub>2</sub> x SP x PT | 1.3<sub>2,143</sub> | 0.27 |  |  |

\*Subscripts indicate degrees of freedom, <sup>^</sup>data were ln transformed to meet assumptions of normality. All models included growth chamber as a random effect.

We found that legumes had a higher shoot % N than grasses, and this resulted in greater total shoot N content for the legumes in both species pairs (**Fig S2**; **Table S1**). Additionally, the *Macroptilium* & *Chloris* species pair generally had greater total shoot N content than *Desmodium* & *Panicum*, driven by the differences in aboveground biomass rather than shoot N concentrations. *Macroptilium* exhibited increased shoot N content under eCO_2_ in monoculture, however there was no change in N content in the mixed pots that included this species (**Fig 2a**). In contrast, *Desmodium* N content was not significantly affected by eCO_2_ in monoculture, however pot total N content in the mixtures with *Panicum* was reduced under eCO_2_ (**Fig 2b**). Legume contributions to shoot N content in mixed pots also differed between the two species pairs, with *Macroptilium* contributing 72%/67% under aCO_2_ and eCO_2_ and *Desmodium* contributing 24%/18% of the total. Importantly, in both species pairs the shoot N content per individual plant in grasses grown with legumes increased (*Chloris*: 51%/83%, *Panicum*: 135%/84%; aCO_2_/eCO_2_, respectively). At the same time, the corresponding shoot N content per individual in the mixed pot legumes declined compared with monoculture pots (*Macroptilium*: -25%/-43%, *Desmodium*: -59%/-85%). For the *Desmodium* & *Panicum* pair this change was large enough that the *Panicum* grown with *Desmodium* contained comparable shoot N content to the *Desmodium* grown in monoculture. Differences in shoot N content among individual species between monoculture and mixed pots (based on ‘PlantType’) are shown in **Table S1** and **Figure S3**.

### N uptake source partitioning: soil N uptake

Despite variability in shoot N contents, the portion of pot total shoot N derived from soil nutrients (**Fig 3**) remained similar among pot types within each species pair (**Table 2**). There were no effects of eCO_2_ on soil-derived shoot N for either species pair, however the fraction of soil-derived N in aboveground plant material in mixed pots was not equally divided between grasses and legumes. Instead, grasses dominated soil nutrient uptake, accounting for 74%/83% for aCO_2_ and eCO_2_ pots in the *Macroptilium* & *Chloris* species pair and 94%/95% in *Desmodium* & *Panicum*. Raw values for δ^15^N for each pot type and CO_2_ treatment are included in **Table S2**. Resin extractable soil nutrients indicated that eCO_2_ did not impact availability (**Fig S4**) of NO_3_^-^, NH_4_^+^ or PO_4_^-3^ (*p* > 0.05). We did, however, find differences in the soil availability of NO_3_^-^ and NH_4_^+^ among pot types within the *Macroptilium* & *Chloris* species pair. *Chloris* monoculture pots had less NO_3_^-^ than in mixed pots (**Fig S4a**), and *Macroptilium* monoculture pots had less NH_4_^+^ than mixed pots (**Fig S4c**).

The ability of plants to take up soil nutrients is contingent on belowground traits, and we found that there were key differences in the root biomass associated with eCO_2_ (*Macroptilium* & *Chloris* **Fig S5a**; **Table S3**) and among pot types (legume/grass/mixed, *Desmodium* & *Panicum*; **Fig S5b**). In general, there was increased root biomass in pots under eCO_2_, driven by significant increases in *Macroptilium* roots in monoculture and for mixed roots in the *Macroptilium* & *Chloris* pots (**Fig S5a**). In the *Desmodium* & *Panicum* species pair there was no impact of eCO_2_, however *Panicum* had the greatest root biomass, followed by the mixed pots and then *Desmodium* in monoculture (**Fig S5b**). Despite these changes in belowground biomass, there were no shifts in the root mass fraction between species pairs, among pot types or associated with eCO_2_ (**Fig S5c** & **d**; **Table S3**).

### N uptake source partitioning: biological nitrogen fixation

We found that a large fraction of the total shoot N was derived from BNF in legume monoculture and mixed species pots, although this differed between legume species and between pot types (**Fig 4**; **Table 3**). *Macroptilium* derived more shoot N via BNF than *Desmodium* (*Macroptilium*: 77/82% of total shoot N in aCO_2_ vs. eCO_2_, *Desmodium*: 32/39%; **Table 3**) in monoculture. Pot level shoot N relied more strongly on BNF in mixtures, with *Macroptilium* & *Chloris* pots increasing reliance on BNF to 84/93% under aCO_2_/eCO_2_ and *Desmodium* & *Panicum* shoot N increasing to 72/48% (**Fig S3b** &**d; Table S1**). Importantly, although both legumes derived less shoot N from BNF in mixture than monoculture (**Fig 4a** & **c**), BNF contributed a greater percentage of the total shoot N in contrast to soil N (**Fig 4d**) for *Desmodium* especially under aCO_2_. Across both species pairs there were low levels of BNF-derived N uptake by grasses (*Chloris*: 3/1%, *Panicum*: 6/1%). However, by mass, the BNF derived N accounted for 29% of the shoot N in *Panicum* in mixtures under aCO_2_, but only 2% under eCO_2_ (**Fig 4b** & **d**).

**Table 3.**
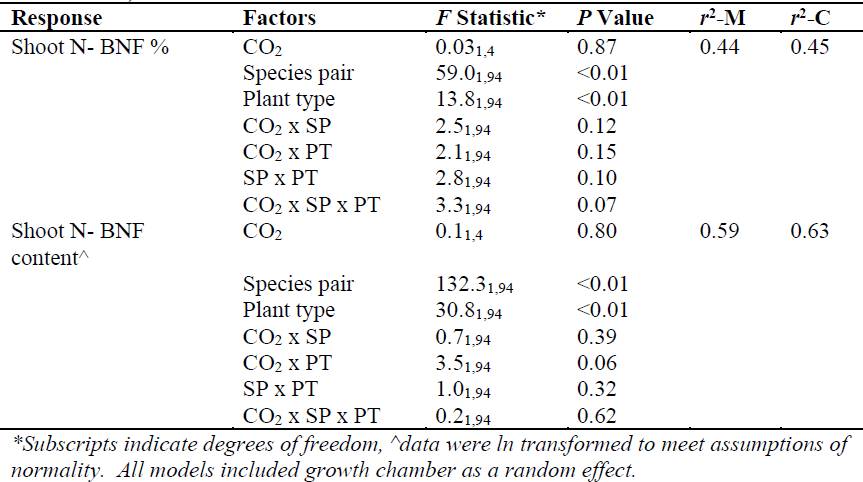
Effects of CO_2_ treatment (CO_2_; ambient, elevated), species pair (SP; *Macroptilium*/*Chloris* and *Desmodium*/*Panicum*) and pot type (PT; legume and mixed-species only), for pot level percent of N derived from biologically fixed N (Shoot N-BNF %; legume and mixed plant types only) and total shoot N derived from BNF (Shoot N-BNF content).

The ability of legumes to support rhizobia in nodules is the main constraint on the potential role of BNF as a source for shoot N, and we found clear differences between species in terms of root nodules (**Fig 5**; **Table 4**). In parallel with total shoot N content, *Macroptilium* had greater nodule biomass (**Fig 5a**) and number of nodules (**Fig 5b**), compared to *Desmodium,* in monoculture. Additionally, both legumes had a significant reduction in nodule biomass and number of nodules when grown with grass, and there were no impacts of eCO_2_ on these measurements. Despite the difference in nodule biomass and number between legume species, *Desmodium* had equivalent levels of instantaneous nodule activity to *Macroptilium*, based on production of ethylene as a proxy for the rate of nitrogen fixation following harvest (**Fig 5c**) and in monoculture there was no impact of eCO_2_ for either species. However, for *Desmodium* grown in mixture with *Panicum*, real-time nodule activity significantly increased for individuals grown under eCO_2_ (**Fig 5c**).

**Fig 3.**
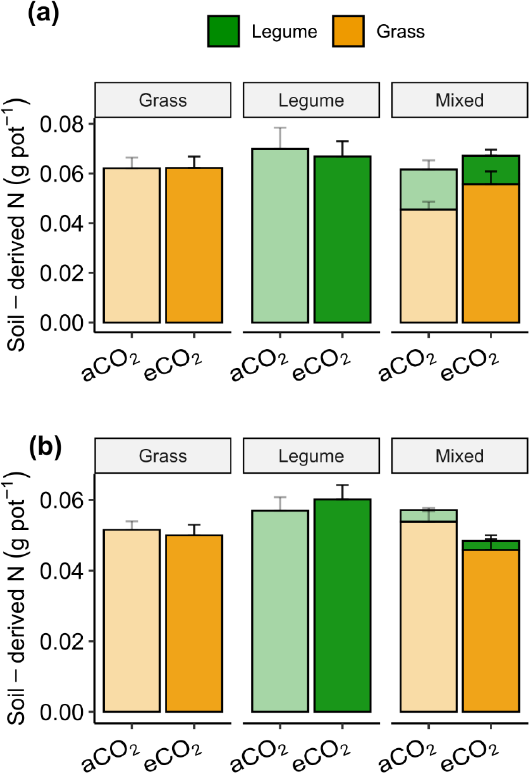
Shoot N content derived from soil for (a) *Macroptilium* & *Chloris* and (b) *Desmodium* & *Panicum* under ambient (aCO_2_) and elevated (eCO_2_) concentrations of CO_2_. Values shown are pot-level means + 1 SE. Note no significant differences with CO_2_ treatment or among pot types were observed.

**Fig. 4.**
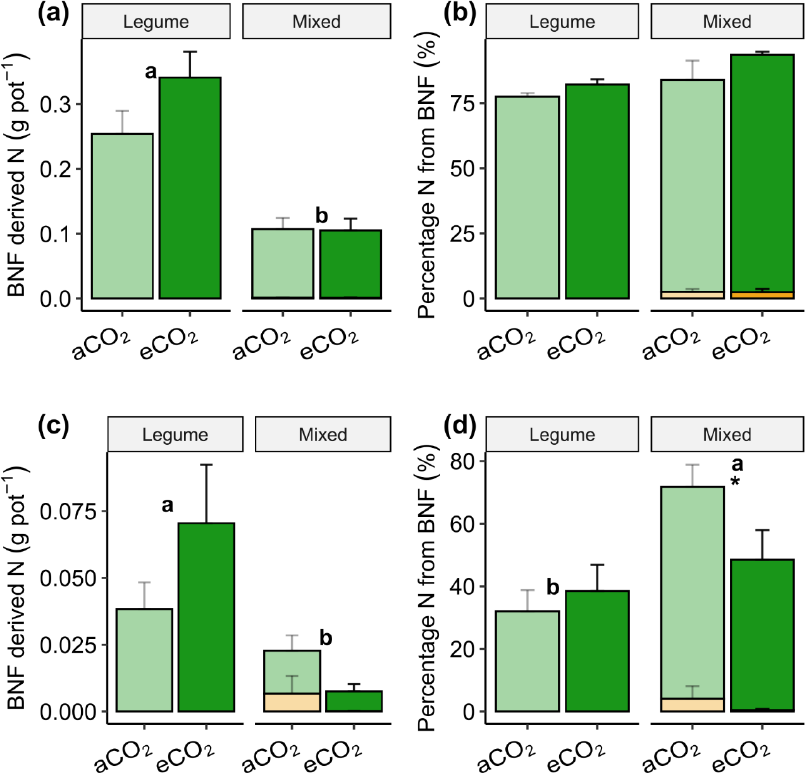
Shoot N content derived from biological N fixation (BNF; a & c) and the proportion of aboveground N content derived from BNF (b & d) for *Macroptilium* & *Chloris* (a & b) and *Desmodium* & *Panicum* (c & d). Values shown are means ± 1 SE. Colours in panels match Fig 1, with green indicating N in legume shoots, and orange indicating N in grass shoots. Notations from Fig. 1.

**Fig 5.**
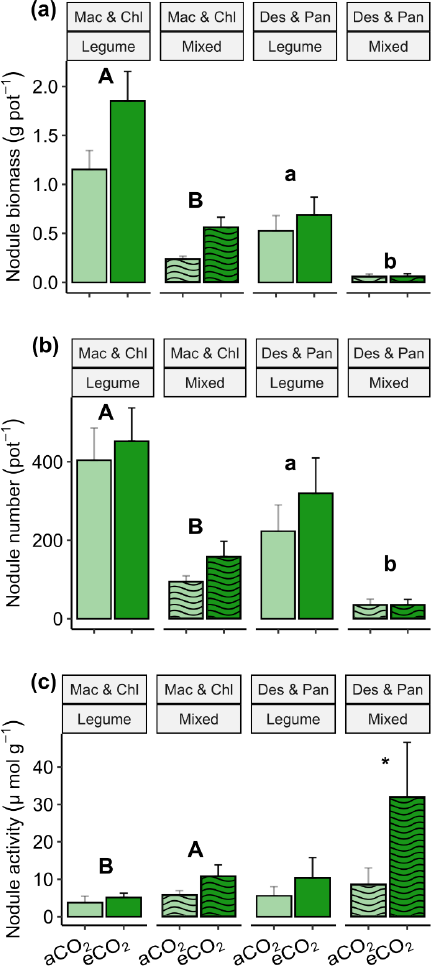
Effects of eCO_2_ on metrics of nitrogen fixation capacity relating to root nodules for legumes grown in monoculture (Legume) and in mixtures with grasses (Mixed; *Macroptilium* and *Chloris*: Mac & Chl, *Desmodium* and *Panicum:* Des & Pan) including (a) the total biomass of nodules per pot, (b) the number of nodules per pot, and (c) root nodule activity as measured by the production of ethylene as a proxy for the rate of nitrogen fixation. Values shown are means ± 1 SE. Differences between plant types within a species pair are indicated by letters (Mac & Chl uppercase, Des & Pan lowercase) and significant differences between CO_2_ treatments are indicated by ‘*’.

**Table 4.** Effects of CO_2_ treatment (CO_2_; ambient, elevated) for pot-level total biomass of root nodules (Nodule biomass), number of root nodules present (Nodule number), and the activity of nodules at the time of harvest (Nodule activity) by species pair (SP; *Macroptilium*/*Chloris* and *Desmodium*/*Panicum*) and plant type (PT; legumes in monoculture vs mixed pots only)

| Response | Factors | F Statistic* | P Value | r<sup>2</sup>-M | r<sup>2</sup>-C |
| --- | --- | --- | --- | --- | --- |
| Nodule biomass<sup>^</sup> | CO<sub>2</sub> | 4.0<sub>1,4</sub> | 0.12 | 0.68 | 0.69 |
|  | Species pair | 65.6<sub>1,68</sub> | <0.01 |  |  |
|  | Plant type | 97.2<sub>1,68</sub> | <0.01 |  |  |
|  | CO<sub>2</sub> x SP | 0.6<sub>1,68</sub> | 0.43 |  |  |
|  | CO<sub>2</sub> x PT | 0.1<sub>1,68</sub> | 0.80 |  |  |
|  | SP x PT | 4.6<sub>1,68</sub> | 0.04 |  |  |
|  | CO<sub>2</sub> x SP x PT | 0.3<sub>1,68</sub> | 0.58 |  |  |
| Nodule number<sup>^</sup> | CO<sub>2</sub> | 0.4<sub>1,4</sub> | 0.55 | 0.55 | 0.58 |
|  | Species pair | 28.8<sub>1,68</sub> | <0.01 |  |  |
|  | Plant type | 72.9<sub>1,68</sub> | <0.01 |  |  |
|  | CO<sub>2</sub> x SP | 0.02<sub>1,68</sub> | 0.88 |  |  |
|  | CO<sub>2</sub> x PT | 0.1<sub>1,68</sub> | 0.8 |  |  |
|  | SP x PT | 4.0<sub>1,68</sub> | 0.05 |  |  |
|  | CO<sub>2</sub> x SP x PT | 0.4<sub>1,68</sub> | 0.52 |  |  |
| Nodule activity<sup>^</sup> | CO<sub>2</sub> | 4.7<sub>1,4</sub> | 0.10 | 0.16 | 0.16 |
|  | Species pair | 0.7<sub>1,68</sub> | 0.42 |  |  |
|  | Plant type | 7.0<sub>1,68</sub> | 0.01 |  |  |
|  | CO<sub>2</sub> x SP | 0.6<sub>1,68</sub> | 0.44 |  |  |
|  | CO<sub>2</sub> x PT | 1.0<sub>1,68</sub> | 0.33 |  |  |
|  | SP x PT | 0.01<sub>1,68</sub> | 0.90 |  |  |
|  | CO<sub>2</sub> x SP x PT | 1.9<sub>1,68</sub> | 0.17 |  |  |

\*Subscripts indicate degrees of freedom, ^data were ln transformed to meet assumptions of normality. All models included growth chamber as a random effect.

### Over-yielding and Complementarity vs. Selection Effects

We found a non-significant positive response of over-yielding for both species’ pairs (**Fig. 6a**). In *Macroptilium* & *Chloris* mixture, over-yielding was mainly determined by the complementarity effect but not the selection effect (**Fig. 6bc**). By contrast, for the *Desmodium* & *Panicum* mixture, the over-yielding was driven more by selection than the complementarity effect (**Fig. 6bc**).

We also found eCO_2_ significantly reduced over-yielding (**Fig. 6a**). In the *Macroptilium* & *Chloris* mixture, complementarity effects shifted from neutral under aCO_2_ to significantly positive under eCO_2_. Meanwhile, selection effects shifted from neutral under aCO_2_ to significantly negative under eCO_2_, meaning that the dominant species had less biomass than expected in the mixture (**Fig. 6c**). In the *Desmodium* & *Panicum* mixture, however, the complementarity effect shifted from significantly positive under aCO_2_ to neutral under eCO_2_, meaning that niche partitioning was weakened. At the same time, the selection effect was unaffected by CO_2_ (**Fig. 6c**), although values were significantly positive under both CO_2_ scenarios, meaning that the dominant species (*Panicum*) consistently had greater biomass in mixtures than proportional production would suggest from monocultures, regardless of CO_2_ treatment. There were no significant effects of eCO_2_ among the two species pairs for either complementarity (CO_2_: F_1,4_ = 0.3, p = 0.62; Species pair: F_1,44_ = 1.9, p = 0.18; CO_2_ x Species pair: F_1,44_ = 2.1, p = 0.15) or selection effects (CO_2_: F_1,4_ = 1.0, p = 0.37; Species pair: F_1,44_ = 94.9, p < 0.01; CO_2_ x Species pair: F_1,44_ = 2.6, p = 0.11). Summing these components of the net biodiversity effect supports the general outcomes of biomass over-yielding only in the *Desmodium* & *Panicum* mixture, with reduced over-yielding under eCO_2_ (**Fig. 6d**). There were, however, no difference between the species pairs in their response to elevated CO_2_ (CO_2_: F_1,4_ = 0.7, p = 0.44; Species pair: F_1,44_ = 3.0, p = 0.09; CO_2_ x Species pair: F_1,44_ = 0.8, p = 0.37).

**Fig. 6.**
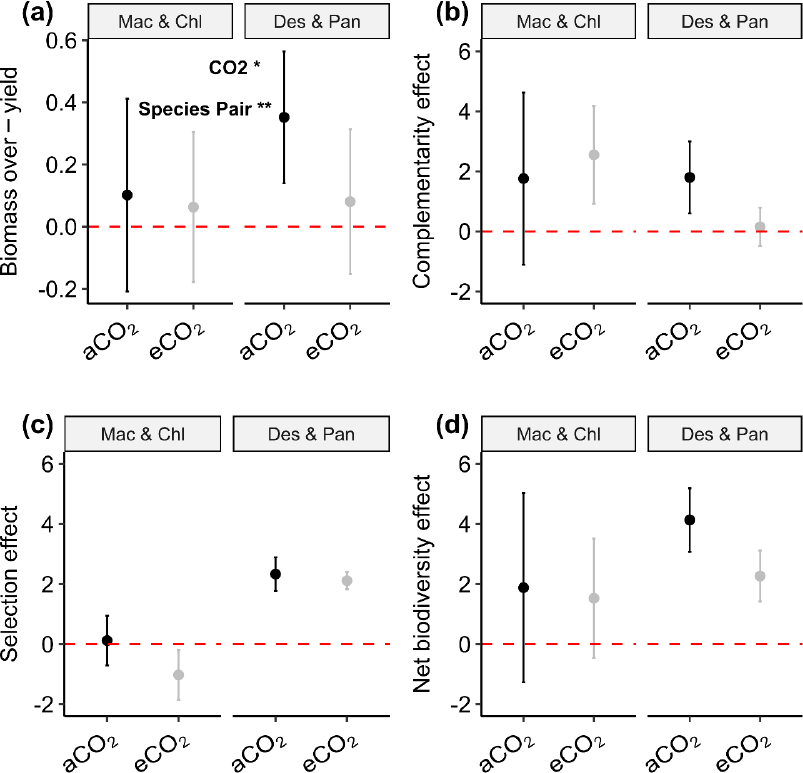
Over-yield effect size for (a) aboveground productivity for pots grown with mixed grass-legume plant types in comparison with standardized summed totals for grasses and legumes grown in monoculture. Species-pair calculations for examining (b) complementarity effect, (c) selection effect and (d) the net biodiversity effect. Positive values indicate an increase relative to monoculture pots (a) or a net positive effect in interaction type between species (b-d). Non-overlapping bars indicate a significant difference from 0. Points shown are means values with 95% CI for *Macroptilium* & *Chloris* (Mac & Chl) mixtures and *Desmodium* & *Panicum* (Des & Pan) mixtures.

## Discussion

In mixed grasslands or pastures containing both grasses and legumes, nutrient niche partitioning among community members can promote species coexistence and ecosystem function, particularly by influencing nutrient availability (Craven et al., 2018; da Silveira Pontes et al., 2015; Fornara & Tilman, 2009). While direct-nutrient sharing is often predicted among community members (Pirhofer-Walzl et al., 2012), nutrient, especially N, niche partitioning is a more common finding in ecological and agricultural studies under field conditions (Nyfeler et al., 2011; Pelzer et al., 2014; Zuppinger-Dingley et al., 2014). In these situations, community members access different pools of available resources, over time and/or space, enabling the persistence of many species by reducing competitive interactions (Adler et al., 2012; Zuppinger-Dingley et al., 2014). Our results found that (1) within tropical grass-legume mixtures, N niche partitioning drove plant-plant interactions such that legumes increased reliance on biological nitrogen fixation (BNF) relative to soil N when grown with grasses leading to increased grass growth per tiller. (2) Elevated CO_2_ (eCO_2_) had varied impact on individual tropical forage species, with benefits in growth for one legume and no direct benefit for grasses, (3) while nodule activity and N content increased for one legume, we found community productivity and N content from tropical legume-grass mixtures were lower under eCO_2_ than aCO_2_, suggesting that legume-grass pairings may benefit ecosystem function to a lesser extent in the face of rising CO_2_; and (4) the species-specific responses associated with over-yielding and responses to eCO_2_, demonstrated differences in complementarity and selection effects operating between the two mixed communities and a lessening of net biodiversity effects under eCO2 associated with reduced niche partitioning between the grass and legume for one species pair.

### Tropical grass-legume mixtures

Relationships between plant functional group diversity and ecosystem function typically predict and find that increased diversity supports greater ecosystem productivity (Hector et al., 1999; Küchenmeister et al., 2012; Loreau & Hector, 2001; Mason et al., 2017; Tilman et al., 2014). In testing this hypothesis, many studies have included grass-forb-legume or grass-shrub-forb combinations experimentally to examine potential over-yielding benefits (Finn et al., 2013), but fewer ecological studies have included C_4_ grasses alongside legumes (Lee et al., 2003; Lilley, Bolger, & Gifford, 2001; Rasmussen et al., 2013). This is despite early results suggesting that the high competitive ability of some C_4_ grasses may functionally contribute to over-yielding (Lambers et al., 2004) and the economic importance of such mixtures in many agricultural grasslands around the world (Butler et al., 2013; Duchene et al., 2017). Previous results have shown that, at a global scale, tropical grasses show lower productivity benefits from the addition of legumes, relative to temperate mixed-species systems (Ashworth et al., 2018). Therefore, increased understanding of the mechanisms that may promote over-yielding are of broad interest, especially in tropical systems. Our study found strong evidence for productivity gains by the grass component in tropical grass-legume mixtures (per tiller), accompanied by declines in the legume partner, which translated into minimal changes at the pot or sward level over a short-term glasshouse experiment. This result aligns with earlier tropical mixed pasture trials that concluded that the relative abundance of grasses and legumes has an important impact on the transfer of N and sward productivity benefits of mixtures, compared to monocultures of the component species (Baba et al., 2011).

Increased productivity in mixed species grasslands, relative to monocultures, is primarily attributed to increased access to resources, including light, water or nutrient availability in ways that promote facilitation. This facilitation can include mechanisms of niche partitioning such that different species access different resource pools in time or space to minimize competition (da Silveira Pontes et al., 2015; Jumpponen et al., 2002; Thilakarathna et al., 2016). Biological or environmental conditions under which different mechanisms promoting nutrient facilitation may dominate are still an area of active research (Brooker et al., 2015), and our study addressed the potential for niche partitioning vs. direct facilitation for N between co-occurring grasses and legumes (Ajayi et al., 2008; Baba et al., 2011; Bell et al., 2017; Miegoue et al., 2016). Despite only limited gains in shoot N for mixed pots, relative to the grass monocultures over our short-term species interactions, this indicated niche partitioning as the main mechanism promoting plant N access between the component species. Under these conditions, the proportion of shoot N derived from BNF increased for legumes even as the total mass of shoot N declined in comparison with monoculture pots, as evidenced by one of our species pairs (*Desmodium*-*Panicum*). In agricultural-grassland settings mixtures of grasses and legumes are often used to specifically improve forage quality and production in the absence of fertilizer applications (Brooker et al., 2016; Li et al., 2015; Warwick et al., 2016). This practice is predicted to play an important role in grassland management for grazed systems as C_4_ grasses become more dominant under changing climate (Butler et al., 2013; McKeon et al., 2009; Still et al., 2019).

There are a variety of mechanisms that enable N niche complementarity in mixed-species grasslands, and it is worth noting that the duration of species associations may play a role in the potential for direct-facilitation of BNF-derived N between for companion grasses in mixed grasslands (Heichel & Henjum, 1991; Thilakarathna et al., 2016). Indeed, a time lag may be associated with the decomposition of legume leaf and root litter that can provide sources for increased ecosystem N availability on longer time scales (Cannell & Thornley, 1998; Kohmann et al., 2019). While there was likely root turnover over the course of our glasshouse experiment, and any senesced leaves were retained in pots, these mechanisms are likely more substantial in field settings especially where soil disturbance may promote incorporation of plant materials directly into soils or where legume residues have sufficient time to undergo decomposition (Brooker et al., 2015; Thilakarathna et al., 2016).

Furthermore, under field conditions, legumes often shift to BNF as plants mature (Edmeades & Goh, 1984; Wery et al., 1986) and this can impact the potential facilitation provided to companion species (Bell et al., 2017). In our study all plants were harvested immediately prior to the first flowering phenophase, and consequently observed patterns are from first-year perennial plants in a rapid growth phase when individuals may rely more strongly on available soil N pools. Grasses in mixed pots in our study did show some evidence for incorporation of BNF-derived N into aboveground plant tissue (up to 29% of shoot N for *Panicum* grown with *Desmodium*), thereby providing evidence of direct facilitation under ambient conditions within this species pair. Additionally, this direct contribution is predicted to increase as swards mature (Thilakarathna et al., 2016).

### Consequences of elevated CO_2_

Previous work has shown that eCO_2_ can increase grassland and crop productivity (Shaw et al., 2002; Terrer et al., 2021), or have no impact over differing time scales, and can result in shifts in the abundance of component species (Bloor et al., 2010; Carroll et al., 2003; Mueller et al., 2016). Under well-watered conditions, C_4_ grasses are not predicted to benefit directly from eCO_2_ and our results follow those predictions (Ainsworth & Long, 2005; Soussana & Lüscher, 2007; Wang *et al*., 2022; but see Reich *et al*., 2018). On the other hand, we found some limited effects of eCO_2_ on legume production, with increased growth in *Macroptilium* likely due to increases in shoot N and root biomass. In contrast, *Desmodium* did not respond to eCO_2_, potentially due to a greater reliance on soil N rather than BNF in monoculture, that may have induced stoichiometric constraints in responding to eCO_2_. This result is in line with research showing that the strongest predictor for an individual’s or species’ positive response to eCO_2_ in legumes is the ability to form nodules (Cernusak et al., 2011; Parvin et al., 2020). For example, *Macroptilium* responded positively to eCO_2_ and also had greater nodule production than *Desmodium*, although the number of nodules was unaffected by eCO_2_. Indeed, despite the positive effect of eCO_2_ on *Desmodium* nodule activity in mixtures, low nodule numbers and biomass overall contributed to a proportional reduction in legume biomass within the mixed sward.

At the sward level, despite some gain in productivity for legume biomass and total shoot N in monoculture (*Macroptilium* only), we didn’t find an increase in productivity for mixed pots under eCO_2_. While this result is in contrast to C_3_ grass-legume mixtures for temperate pastures in Australia where eCO_2_ has been shown to increase total shoot N yield (Lilley, Bolger, Peoples, et al., 2001), it is consistent with evidence for limited responses to eCO_4_ by C_4_ grasses and legumes. One explanation may be related to shifts in the nutritional quality of component species under eCO_2_, as reductions in quality for aboveground biomass have been reported across ecosystems (Augustine et al., 2018; Bhargava & Mitra, 2021). Our species experienced limited shifts in quality based on shoot N%, including lower N in *Macroptilium* when grown with *Chloris* and in *Panicum* when grown with *Desmodium*. Even so, these patterns align with other studies measuring nutritional shifts in both tropical grasses and legumes under low soil nutrients where legume shoot N% declined in mixture under eCO_2_ (Edwards et al., 2006).

While the use of confined soil space to test mechanisms for the source of nutrient use and acquisition between grasses and legumes is typical in glasshouse settings, there are some key factors that are likely to impact the species interactions and dynamics in response to eCO_2_ that are not captured here that are important for overarching messages in relation to nutrient facilitation. Our study found that eCO_2_ conditions differentially altered the shoot N% between legumes in mixture but had no impact on legumes grown in monoculture. Under field settings, the effects of herbivory on legumes that maintain a higher nutritive quality relative to surrounding species may result in a comparative loss in abundance or persistence within the community, ultimately reducing the potential over-yield or net biodiversity effect (Rogers et al., 2009). Additionally, literature reviews on field studies have generally found an increase in root length and biomass associated with eCO_2_ in grasslands (Dieleman et al., 2012; Nie et al., 2013), a pattern that was seen in pots containing *Macroptilium* in this study. While our pots were not root-bound by the end of the experiment, other key differences between field and glasshouse conditions may have limited further responses, for example Nie et al. (2013) also found a shift in the depth distribution of roots that is not possible under non-field settings. Such spatial re-distribution of roots with depth can play a major role in complementarity effects between species within mixtures (Oram et al., 2018). Finally, field conditions also typically introduce variation in other resources that modify the interactions between eCO_2_ and nutrient-use, in particular water availability. Typically, C_4_ grasses have been shown to benefit from eCO_2_ under drier soil conditions, further altering the potential interactions between grasses and legumes in mixed swards (Ainsworth & Long, 2005; Reich *et al*., 2014, but see Reich *et al*., 2018).

### Over-yield implications under future CO_2_ scenarios

Reliance on different sources of N among species within mixed plant communities can ultimately reduce competition especially under low nutrient availability (Ball et al., 2021; Elias & Agrawal, 2021). The increased N niche partitioning between grasses and legumes grown in mixture found in this study support this conclusion. Furthermore, the species pair that had a substantial shift in N source use between monoculture and mixtures was also associated with a significant over-yielding benefit. This suggests that greater plasticity in niche space may promote species coexistence and enhance ecosystem production. Belowground, the idea of nutrient form (nitrate vs. ammonium) niche plasticity supporting coexistence has been well-established for cold-climate grasslands (Ashton et al., 2010) and aboveground light-use plasticity has been shown to translate into increased production in experimental grasslands (Meilhac et al., 2020).

Importantly, however, we found that eCO_2_ reduced over-yielding for both aboveground biomass and total shoot N in our tropical grass-legume mixtures. While there were no statistical effects of eCO_2_ on complementarity and selection effects directly, we found key differences in the relative importance of these effects in contributing to the over-yield/net biodiversity effect, with positive complementarity and selection effects together resulting in increased sward level over-yielding that was greatest under aCO_2_.

## Conclusions

Plant responses to eCO_2_ are largely dependent on soil water and nutrient availability. Therefore, shifts in plant-plant interactions including competition (evidenced by niche partitioning) or facilitation among community members may drive diverse ecosystem responses to changes in CO_2_ concentrations. Among our tropical pasture species, we found that legumes grown in monoculture relied approximately equally on available soil N pools and BNF for their N requirements. When grown with tropical grasses, however, legumes shifted to rely either equally or more strongly on BNF, the latter indicative of stronger niche partitioning and less successful competition for soil nutrients with grasses. This separation of niche space was weakened under elevated CO_2_ conditions, ultimately reducing legume production and minimizing over-yielding benefits. Disentangling contributions of niche partitioning and abiotic facilitation, here quantified using stable isotopes, are key to understanding grass-legume contributions to ecosystem function under future climate conditions.

## Acknowledgements

The authors would like to acknowledge individuals whose efforts supported the data collection of this work, including Kathryn Fuller, Alexandra Boyd, Shania Therese Didier Serre, Samantha Weller, Benjamin Capel, Minh Doan. We thank Neha Mohan Babu for comments on the final manuscript. Valuable input from PACE Advisory Board members - Doug McNicholl (Meat and Livestock Australia), Cath Lescun (Dairy Australia), Catherine Phelps (Dairy Australia), Tom Dickson (Heritage Seeds), Brendan Cullen (University Melbourne) and Suzanne Boschman (Department of Primary Industries) - is also gratefully acknowledged. This project was supported by funding from Meat and Livestock Australia’s Donor Company (P.PSH. 0793), Dairy Australia (C100002357) and Western Sydney University. A version of this manuscript was included as a preprint on BioRxiv.

## Author Contributions

Authors ACC, HZ, and GWK designed the experiment with input from SAP, EP, BM, JRP, JMP, and KLMC. ACC, HZ, GWK, and KLMC performed experiments and conducted glasshouse work. ACC conducted statistical analyses with input from HZ, JRP and FI; ACC and HZ wrote the paper with input from all co-authors on draft iterations.

## Data Availability

The data that support the findings of this study are openly available in Dryad at 10.5061/dryad.nk98sf7ww.

