## Supplemental Materials for "Nitrogen niche partitioning between tropical legumes and grasses conditionally weakens under elevated CO_2_"

### Supporting Information

**Table S1**) Effects of $CO_2$ treatment ($CO_2$; ambient, elevated), species pair (SP; *Macroptilium*/*Chloris* and *Desmodium*/*Panicum*) and plant type (PT; grass, legume, grass grown with legume: L-grass, and legume grown with grass: G-legume), for aboveground plant percent N (%), shoot N content (g) and BNF-derived shoot N (g N $g^{-1}$ dry mass).
.

| **Response** | **Factors** | ***F* Statistic*** | ***P* Value** | **$r^2$-M** | **$r^2$-C** |
|---|---|---|---|---|---|
| Shoot N %^ | $CO_2$ | $4.9_{1,4}$ | 0.09 | 0.84 | 0.85 |
| | Species pair | $30.1_{1,126}$ | <0.01 | | |
| | Plant type | $369.3_{3,139}$ | <0.01 | | |
| | $CO_2$ x SP | $0.1_{1,126}$ | 0.80 | | |
| | $CO_2$ x PT | $0.7_{3,139}$ | 0.54 | | |
| | SP x PT | $9.0_{3,139}$ | <0.01 | | |
| | $CO_2$ x SP x PT | $2.3_{3,139}$ | 0.08 | | |
| Shoot N Content^ | $CO_2$ | $0.4_{1,4}$ | 0.56 | 0.70 | 0.70 |
| | Species pair | $112.4_{1,191}$ | <0.01 | | |
| | Plant type | $81.6_{3,191}$ | <0.01 | | |
| | $CO_2$ x SP | $3.9_{1,191}$ | 0.05 | | |
| | $CO_2$ x PT | $3.6_{3,191}$ | 0.01 | | |
| | SP x PT | $39.4_{3,191}$ | <0.01 | | |
| | $CO_2$ x SP x PT | $0.8_{3,191}$ | 0.51 | | |
| BNF-derived shoot N | $CO_2$ | $1.2_{1,4}$ | 0.34 | 0.81 | 0.81 |
| | Species pair | $32.9_{3,191}$ | <0.01 | | |
| | Plant type | $279.0_{3,191}$ | <0.01 | | |
| | $CO_2$ x SP | $0.6_{1,191}$ | 0.44 | | |
| | $CO_2$ x PT | $1.0_{3,191}$ | 0.40 | | |
| | SP x PT | $10.2_{3,191}$ | <0.01 | | |
| | $CO_2$ x SP x PT | $1.3_{3,191}$ | 0.28 | | |

**Subscripts indicate degrees of freedom, ^data were ln transformed to meet assumptions of normality. All models included growth chamber as a random effect.*

**Table S2**) Mean $\delta^{15}N$ (± se) for dried shoot biomass among tropical pasture species under ambient $CO_2$ ($aCO_2$) and elevated $CO_2$ ($eCO_2$) treatments for individuals grown in monoculture vs. mixed pots

| **Species** | **$CO_2$ Treatment** | **Monoculture** | **Mixed** |
|---|---|---|---|
| *Chloris* | $aCO_2$ | 3.93 ± 0.18 | 4.17 ± 0.20 |
| | $eCO_2$ | 3.74 ± 0.22 | 3.82 ± 0.15 |
| *Desmodium* | $aCO_2$ | 2.36 ± 0.48 | -0.01 ± 0.44 |
| | $eCO_2$ | 1.81 ± 0.52 | 1.25 ± 0.60 |
| *Macroptilium* | $aCO_2$ | -1.20 ± 0.09 | -1.31 ± 0.64 |
| | $eCO_2$ | -1.54 ± 0.13 | -2.12 ± 0.08 |
| *Panicum* | $aCO_2$ | 4.90 ± 0.16 | 4.44 ± 0.31 |
| | $eCO_2$ | 4.40 ± 0.10 | 4.64 ± 0.11 |

**Table S3**) Effects of $CO_2$ treatment ($CO_2$; ambient, elevated), pot type (PT; grass, legume, mixed-species), for pot level total root biomass and root mass fraction (RMF; root biomass standardized by total biomass)

| **Response** | **Factors** | **F Statistic** | **P Value** | **$R^2M$** | **$R^2C$** |
|---|---|---|---|---|---|
| Root biomass | $CO_2$ | $9.2_{1,4}$ | 0.04 | 0.43 | 0.43 |
| | Species pair | $48.0_{1,110}$ | <0.01 | | |
| | Plant type | $9.5_{2,110}$ | <0.01 | | |
| | $CO_2$ x SP | $1.9_{1,110}$ | 0.18 | | |
| | $CO_2$ x PT | $1.6_{2,110}$ | 0.20 | | |
| | SP x PT | $5.2_{2,110}$ | <0.01 | | |
| | $CO_2$ x SP x PT | $1.3_{2,110}$ | 0.29 | | |
| RMF | $CO_2$ | $0.3_{1,4}$ | 0.61 | 0.05 | 0.14 |
| | Species pair | $0.0_{1,107}$ | 0.97 | | |
| | Plant type | $2.0_{2,107}$ | 0.14 | | |
| | $CO_2$ x SP | $0.7_{1,107}$ | 0.39 | | |
| | $CO_2$ x PT | $0.3_{2,107}$ | 0.72 | | |
| | SP x PT | $0.7_{2,107}$ | 0.51 | | |
| | $CO_2$ x SP x PT | $0.0_{2,107}$ | 0.99 | | |

*All models included growth chamber as a random effect.*

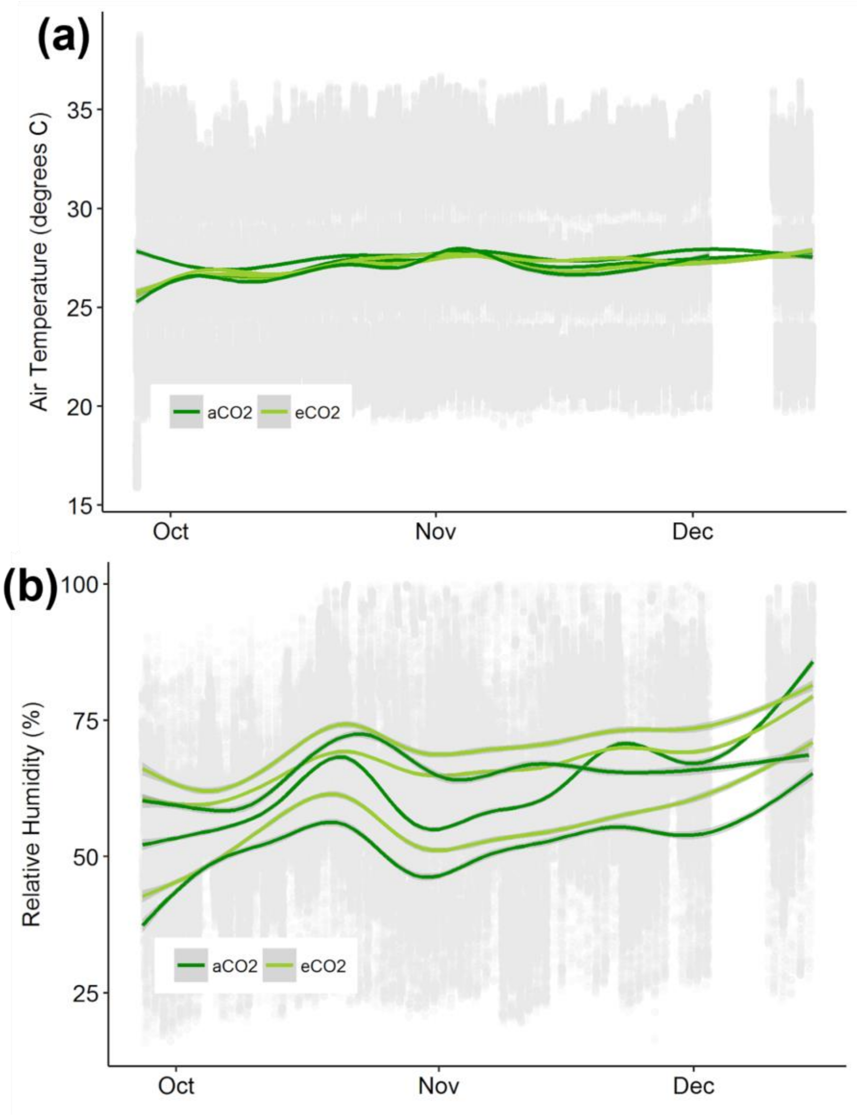


**Fig S1**. Chamber environmental conditions throughout experiment including A) average chamber air temperature and B) air relative humidity. Temperature followed a diurnal regime (00-06 hours: 22°C, 06-09 hrs: 27°C, 09-18 hrs: 32°C, 18-21 hrs: 27°C, 21-00 hrs: 22°C) set to match spring-summer temperatures in regions where tropical pastures are grown in southern Queensland and northern New South Wales, Australia.

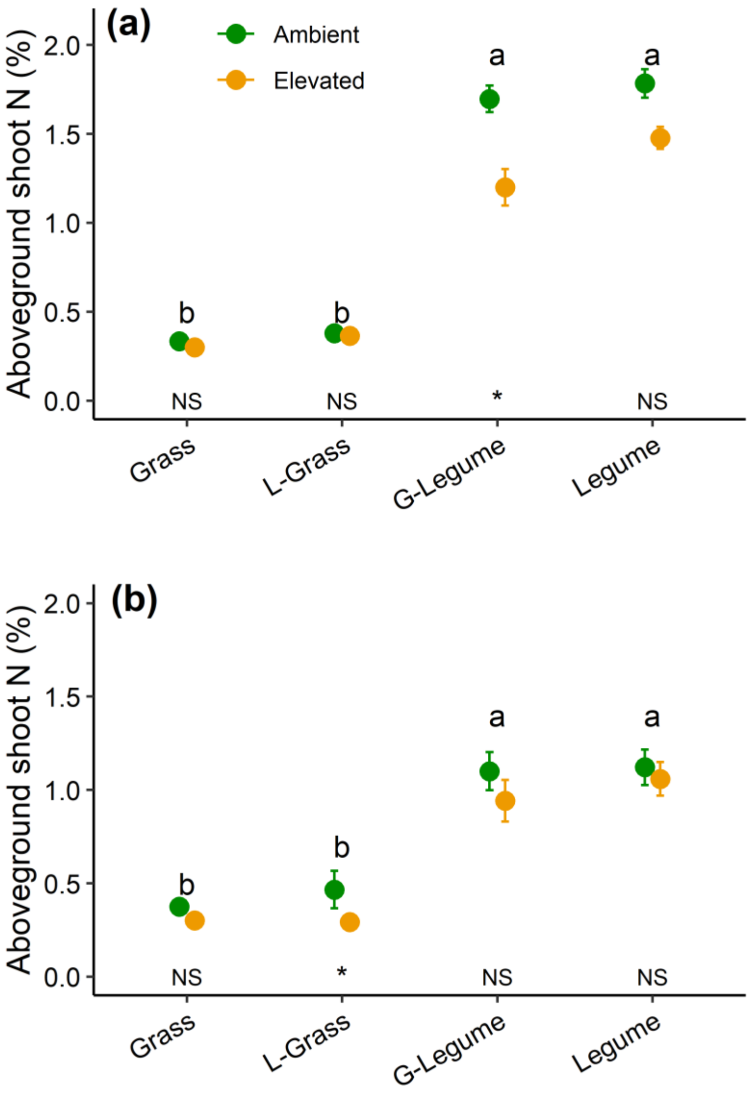


**Fig S2.** Shoot % N for *Macroptilium* & *Chloris* (a) and *Desmodium* & *Panicum* (b). Values shown are means ± 1 SE. There were no significant differences between $CO_2$ treatments, while significant differences among plant types (grass, legume, grass grown with a legume: L-Grass, and legume grown with a grass: G-Legume) are indicated by differing letter designations.

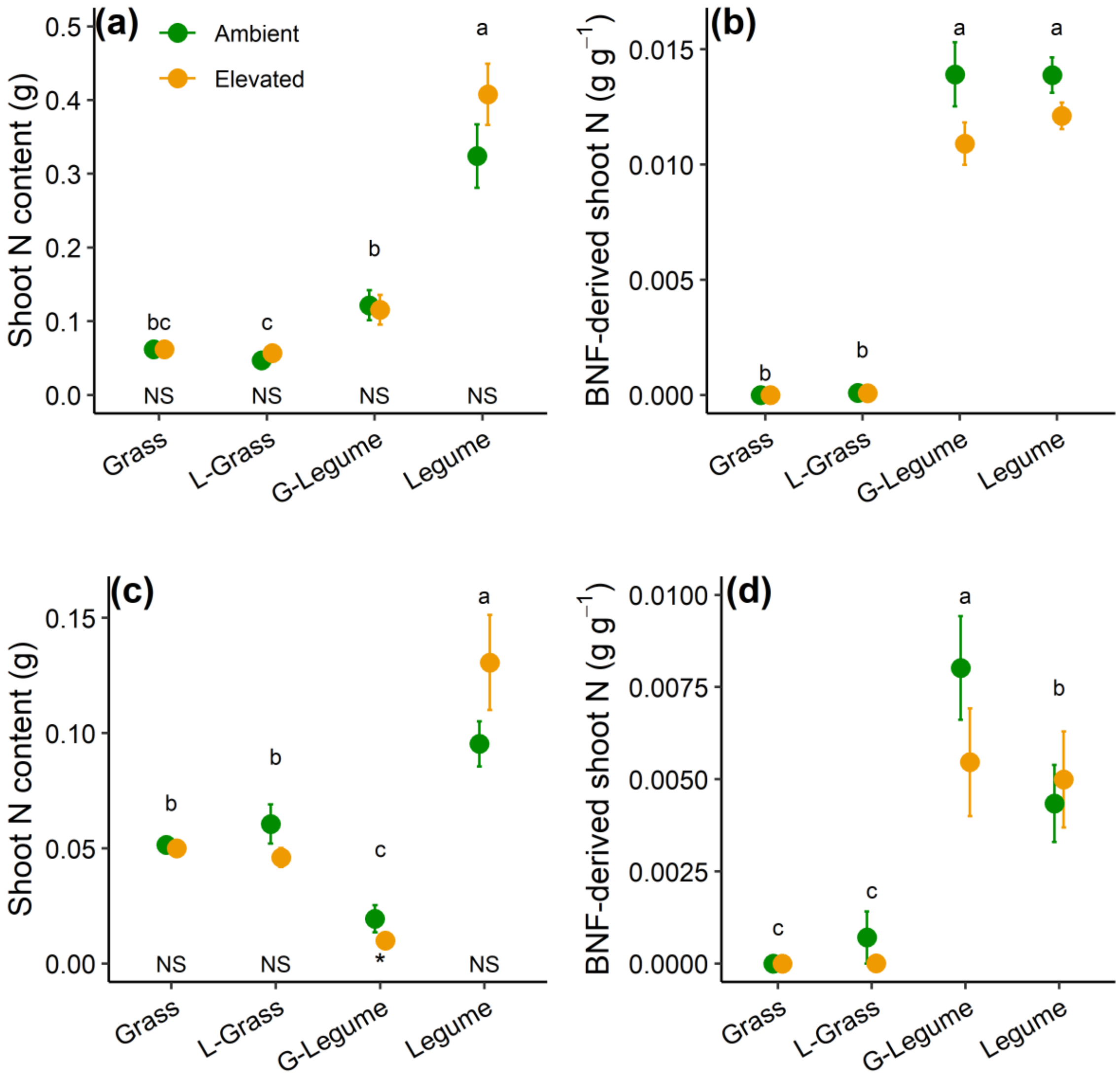


**Fig S3.** Shoot N content (g; a & c) and BNF-derived shoot N (g N $g^{-1}$ dry mass; b &d) for *Macroptilium* & *Chloris* (a & b) and *Desmodium* & *Panicum* (c & d). Values shown are means ± 1 SE. Significant differences ($p < 0.05$) between $CO_2$ treatments are indicated by '*' for shoot N, and significant differences among plant types (grass, legume, grass grown with a legume: L-Grass, and legume grown with a grass: G-Legume) are indicated by differing letter designations.

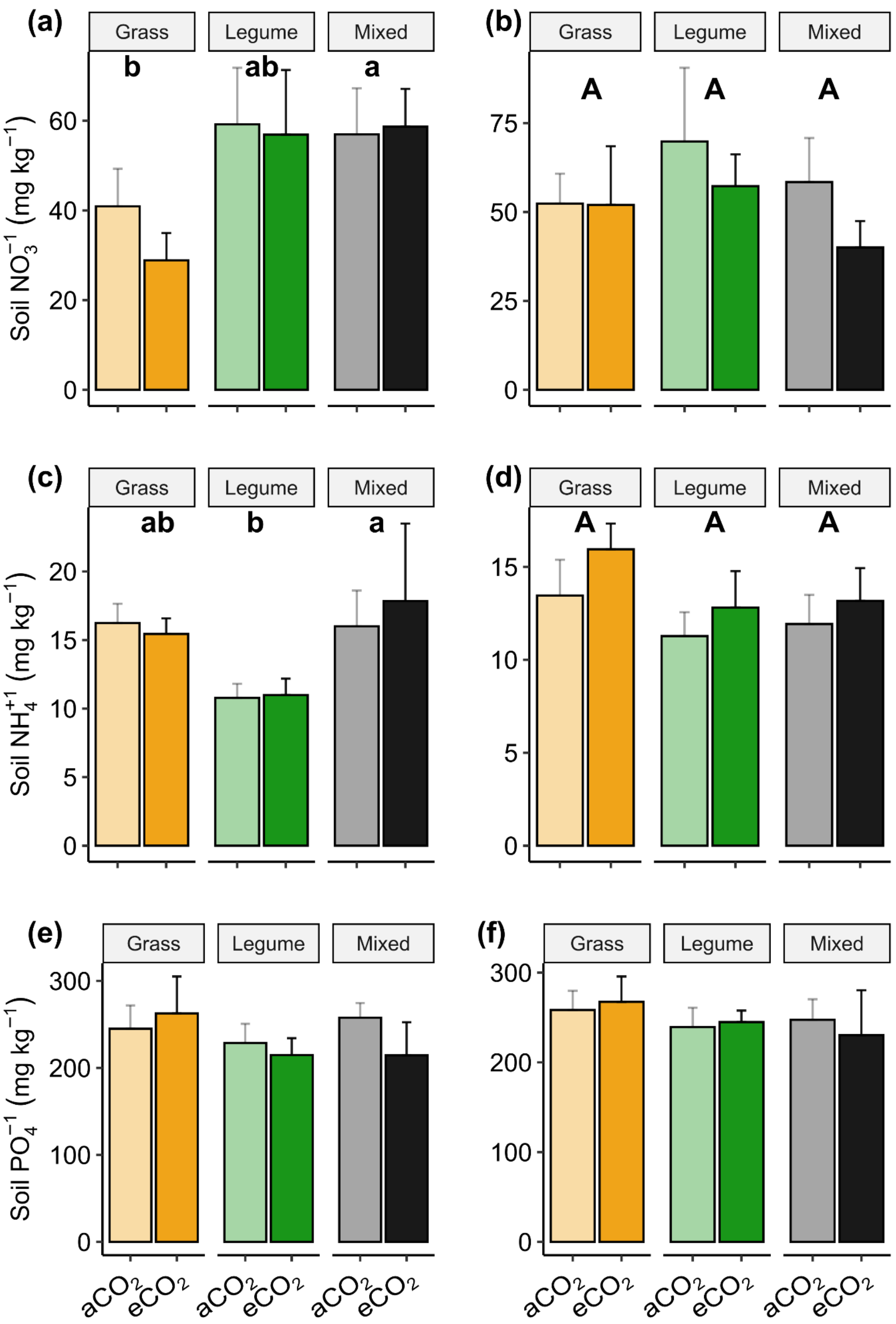


**Fig S4)** Effects of elevated $CO_2$ on resin extractable soil nutrients including $NO_3^-$ (a, b), $NH_4^+$ (c, d) and $PO_4^-$ (e, f) for tropical pasture species grown independently (Grass and Legume in legend) and together (Mixed) under ambient ($aCO_2$) and elevated $CO_2$ ($eCO_2$) conditions for (a & c) *Macroptilium* and *Chloris* and (b & d) *Desmodium* and *Panicum* species pairs. Bars indicate treatment means ± 1 standard error and same letters are used to indicate non-significant differences among pot types for a species pair.

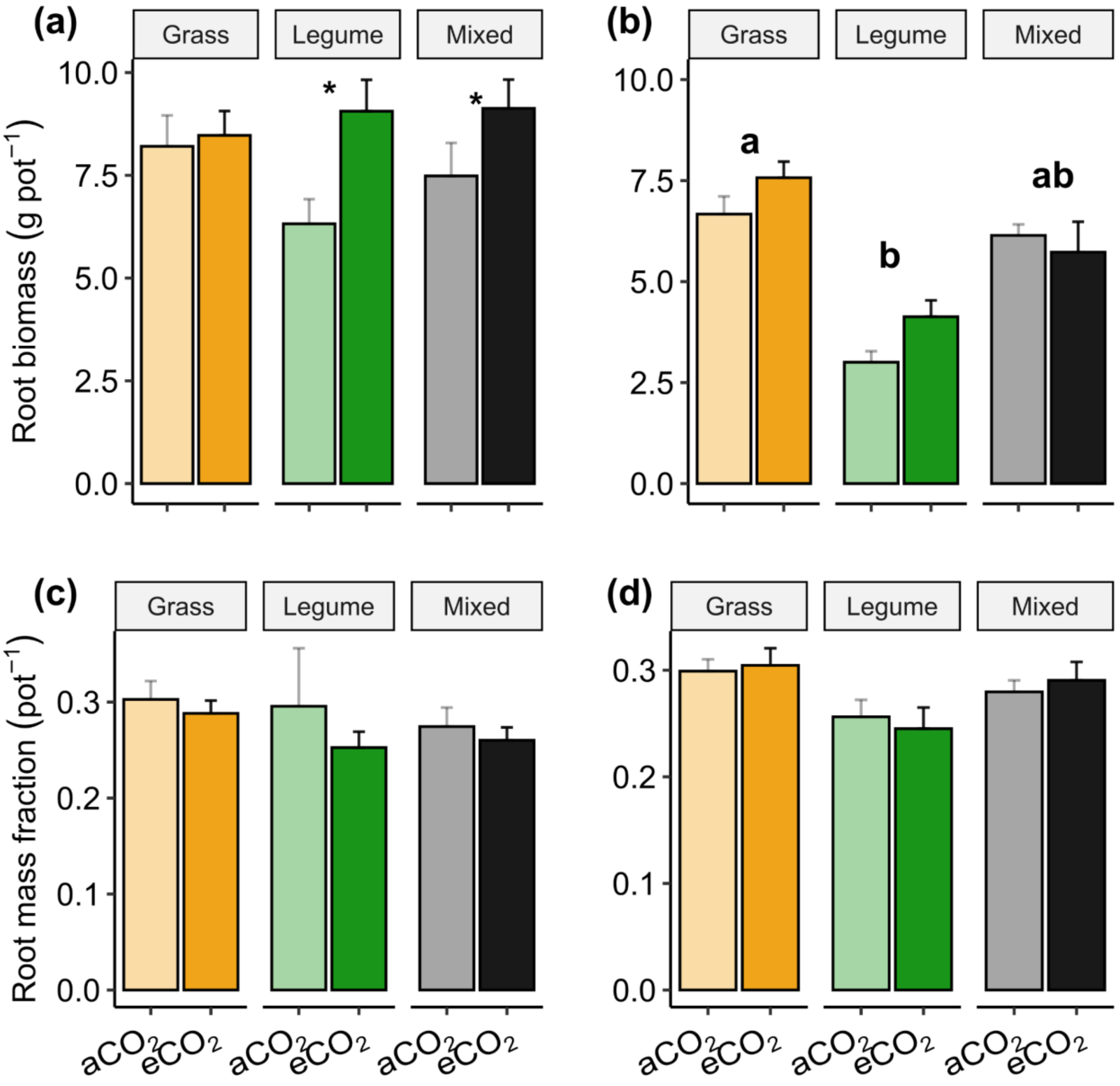


**Fig S5)** Root biomass (a & b) and root mass fraction (c & d) for *Macroptilium*/*Chloris* (a & c) and *Desmodium*/*Panicum* (b & d) pots exposed to $CO_2$ treatment for plants grown in monoculture and in mixed species pots. Bars indicate mean values, plus standard error. Significant differences ($p < 0.05$) between $aCO_2$ and $eCO_2$ treatments are indicated by '*', and significant differences in total pot biomass among plant types (grass, legume, mixed) are indicated by differing letter designations.
